# Observing the self and other in motion modulates the excitability of vestibulocollic reflexes

**DOI:** 10.1101/2022.09.16.503320

**Authors:** Estelle Nakul, Diane Deroualle, Marion Montava, Jean-Pierre Lavieille, Christophe Lopez

## Abstract

Vestibular inputs from the inner ear are at the basis of the vestibulo-spinal and vestibulocollic reflexes involved in balance control. Studies have focused on how attentional load and emotions influence balance, but low-level social cues, such as observing human bodies in motion, have been neglected. Yet, individuals observing another person in a challenging posture or in motion can experience imbalance, indicating that sensorimotor resonance between self and others is involved. The present study examines how the observation of videos depicting human bodies in motion modulates well-established neurophysiological signatures of vestibular information processing. The excitability of vestibulocollic reflexes was assessed by analyzing the waveform of vestibular-evoked myogenic potentials (VEMPs) over the sternocleidomastoid and trapezius muscles of 25 healthy participants (13 females, 12 males). Here we show that observing human bodies undergoing passive whole-body rotations reduced the VEMPs amplitude when compared to observing an object. Importantly, the modulation depended on the person depicted in the video as VEMPs were reduced when observing oneself, compared to someone else being moved. Direction-specific effects and electromyography recordings ruled out non-specific emotional and attentional effects. These results show that the vestibular system is sensitive to observing human bodies in motion, establishing new connections between social neuroscience and vestibular neurophysiology.

**Significance Statement:** Vestibulocollic reflexes are thought to be consistent and of short latency. Yet, previous results show that observing conspecifics influences balance. We combined approaches from social neuroscience and vestibular electrophysiology to describe how the observation of self and other bodies in motion influences vestibular information processing. The results show that observing human bodies in motion reduces the amplitude of vestibulocollic reflexes involved in the stabilization of the head and balance. These results establish new relations between the sense of balance and social cognition and challenge classical views in vestibular neuroscience.

## Introduction

Vestibular signals originating from the inner ear are essential for multisensory self-motion perception and accurate balance control^1^. These signals trigger stabilizing reflexes in postural and neck muscles when the body is translated or rotated, and damage to the inner ear impairs body orientation and stabilization^2^. The vestibular control of balance and of head stabilization in space is supported by projections from the vestibular nerve to the vestibular nuclei in the brainstem, and then to alpha and gamma motoneurons^3^. Vestibulo-spinal and vestibulocollic reflexes that maintain balance and stabilize the head in space are consistent and of short latency^4–6^. In humans, vestibulocollic reflexes are now classically studied by recording vestibular-evoked myogenic potentials (VEMPs) over cervical muscles. Cervical VEMPs consist of inhibitory reflexes evoked by auditory clicks^7^ or by electrical impulses over the vestibular nerve^7,8^ and are characterized by a biphasic p13-n23 wave^9^. Although the pattern of excitatory and inhibitory connections between the different vestibular receptors and neck muscles is well described^6,10^, there is scarce description of how cognitive, emotional and environmental factors influence vestibulocollic reflexes^11^.

Recent evidence suggests that self-motion perception and vestibular reflexes are not as immune to emotions and cognition, as it is sometimes posited. For example, postural threats on participants standing on an elevated platform, increase the amplitude of cervical VEMPs^12^. Moreover, participants involved in a cognitive task while standing or walking show a decreased balance^13^, whereas patients with phobic postural vertigo exhibit a balance behavior that is closer to controls when distracted with a cognitive dual task^14^. To date, there has been little research about the effects of the observation of conspecifics on vestibular perception and cognition, as is typical for other sensory systems^15^. It has been shown that observing another person in a challenging posture or in motion may evoke an imbalance in the observer^16^. Furthermore, observing videos of bodies being passively rotated on a motorized chair modulated performance in a self-motion detection task^16,17^. Of note, patients with peripheral vestibular disorders report discomfort and poorer balance control when surrounded by crowds of people moving around them^15,18,19^, but the underpinnings of the influence of other motion observation on vestibular information processing are still unknown.

In the present study, healthy participants observed videos of human bodies or an object undergoing passive whole-body motion while we assessed the excitability of vestibulocollic reflexes by evoking cervical VEMPs with galvanic vestibular stimulation. We hypothesized that observing the passive motion of one’s own body, of another unknown body, or an object, would modulate VEMPs waveforms differently. We also analyzed how empathy traits^20^ related to VEMPs modulation. This was motivated by behavioral data showing that empathy influenced self-motion perception abilities during the observation of other bodies in motion^17^. Personality traits were also shown to modulate brain response to vestibular stimulation^21^.

We found that observing the passive motion of human bodies (self and other) or of an object have different effects on VEMPs, supporting our assumption that vestibular information processing can be modulated by self-other representations, a crucial feature of social cognition.

## Results

### Self-reports and behavioral results

Participants experienced moderate illusory self-motion when observing passive rotations of a body/object (**Figure 1D**). Some participants reported that Self videos evoked “a motion of the head, like in a mirror”, or “the sensation of being rotated”. By contrast, participants did not report self-motion during Other videos (e.g., “I was looking at the other person being rotated, but I did not have the sensation I was rotated”). Friedman’s ANOVA revealed a near-significant main effect of Video on illusory self-motion (χ^2^(2) = 5.49, p = 0.06). Two-sided Wilcoxon signed-rank tests showed marginally higher illusory self-motion for Self than Other videos (Z = 1.83, p = 0.068) and for Self than Object videos (Z = 1.86, p = 0.06), whereas illusory self-motion was similar for Other and Object videos (Z = 0, p = 1). There was no significant effect of Direction of rotation on illusory self-motion (χ^2^(1) = 1.0, p = 0.32).

**Fig 1.**
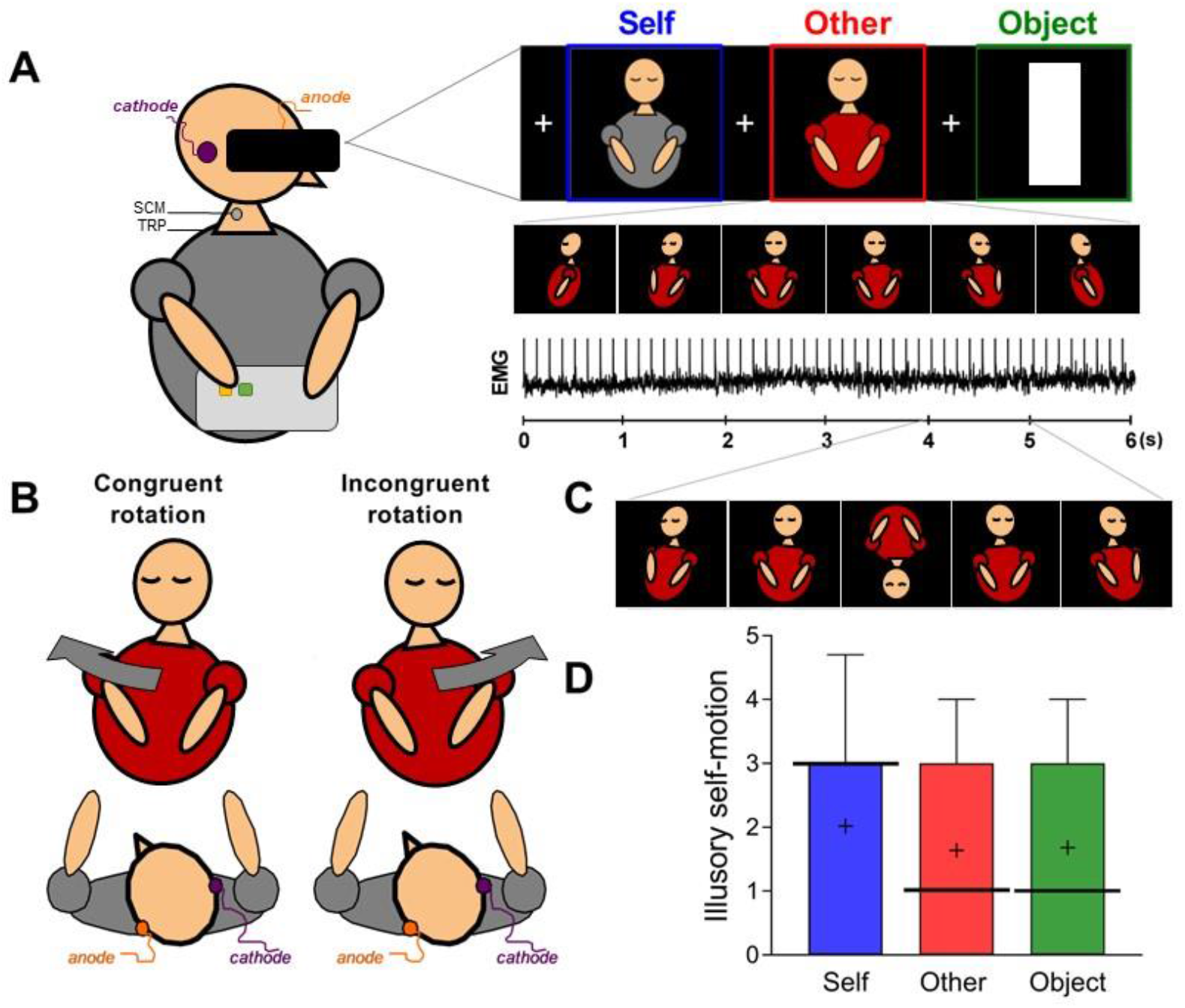
Experimental setup and procedures. **A.** Binaural galvanic vestibular stimulation (GVS) and electromyographic (EMG) recordings recorded over the sternocleidomastoid (SCM) and the trapezius (TRP) muscles. Participants actively maintained their head flexed towards the torso and rotated towards the anode to contract the SCM and the TRP under the cathode. Six-second videos showing the passive rotation of the participant (“Self videos”), an unknown person (“Other videos”), or a white object (“Object videos”) were presented in a head-mounted display. GVS was applied during the videos and electric artefacts were visible on the EMG signal. **B.** “Congruent rotation” showed rotations in the direction matching what participants would have seen of their initial head rotation in a mirror, whereas “Incongruent rotations” showed rotations in the opposite direction (specular congruency). **C.** Example of the vertical inversion of the image (100 ms; 1.5 s, 3 s or 4.5 s after the video onset), present in 25% of the videos. **D.** Box-and-Whisker plots illustrate the intensity (0 = “not at all”; 7 = “absolutely felt something”) of illusory self-motion for each category of Videos. The top and bottom ends of the whisker represent the 90^th^ and 10^th^ percentiles of the distribution, the bold horizontal line represents the median and the black cross represents the mean.

Participants detected the inversion of the images in the videos with a mean accuracy of 98%, indicating that they attended to the task. There was a near-significant effect of the Video on accuracy (χ^2^(2) = 5.75, p = 0.06) and no effect of the Direction of rotation (χ^2^(1) = 0.53, p = 0.47). When exploring the statistical trend of the effect of Video, we found that accuracy did not differ between categories of videos (Self vs. Other: Z = 0.24, p = 0.81; Self vs. Object: Z = 1.7, p = 0.09; Other vs. Object: Z = 1.6, p = 0.11).

### VEMPs waveform analyses

**Figures 2** and **3** illustrate the effect of Video (Friedman’s ANOVAs) on the corrected VEMP amplitude over the SCM and TRP muscles, separately for Congruent and Incongruent rotations. Different patterns of modulation by the Video were found for Congruent and Incongruent rotations. For Congruent rotations, we found a significant main effect of the Video for both SCM and TRP muscles in time windows spanning the p13 wave (SCM muscle: 62 consecutive significant data points [csdp.] at p < 0.05, from 12.1 to 15.0 ms; TRP muscle: 26 csdp., 17.8–19.3 ms), the decreasing portion after the peak of component n23 (SCM: 50 csdp., 23.8–26.8 ms; TRP: 34 csdp., 27.5–29.5 ms), and the decreasing portion after the peak of P2 (SCM: 24 csdp., 40.7–42.1 ms, followed by 65 csdp., 42.4–46.3 ms), as shown by the colored areas superimposed on VEMPs waveforms in **Figures 2A** and **3A**. There was also a main effect of Video spanning the peak of component N2 on the TRP muscle (29 csdp., 56.2–57.9 ms, followed by 39 csdp., 59.3–61.6 ms). By contrast, Friedman’s ANOVAs for the Incongruent rotations yielded no effect of the Video on the VEMPs amplitude during the p13-n23 biphasic wave, and the P2–N2 (**Figures 2B and 3B**). There were only 28 csdp. from 68.1 to 69.8 ms, thus after the later components, for the TRP muscle. Accordingly, we report below only results from post-hoc analyses comparing the waveform of the VEMPs between each category of Videos in the Congruent rotation condition.

**Fig 2.**
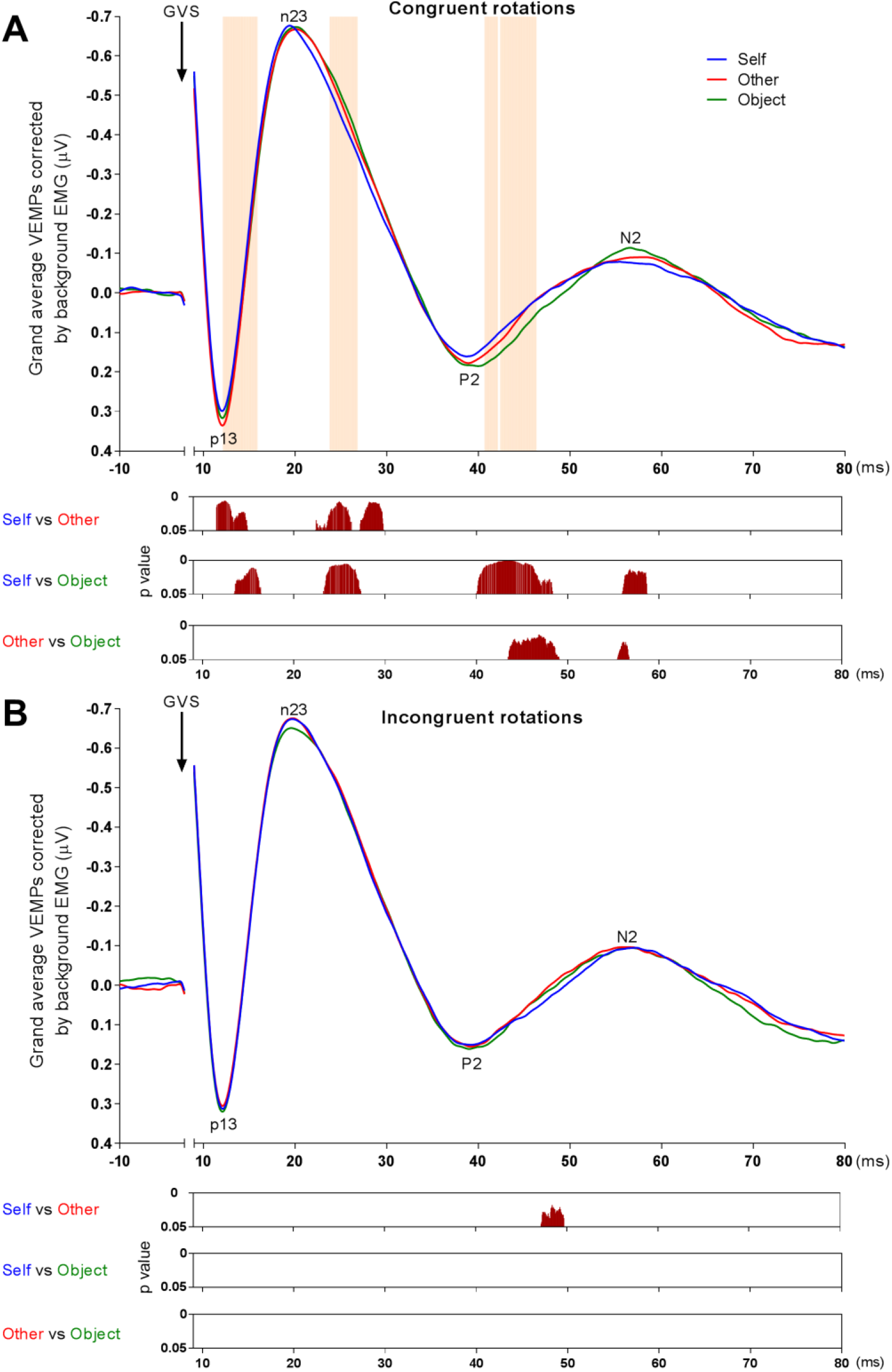
VEMPs recorded over the SCM muscle. Grand-average VEMPs (n = 24) are showed separately for Congruent rotations (A) and Incongruent rotations (B) for each category of Videos. Colored areas in the upper panel show periods with a significant main effect of the Videos (Friedman’s ANOVA). The three lower panels present results from post-hoc analyses (Wilcoxon signed-rank test). Analyses were corrected for temporal autocorrelation by using the constraints of 20 consecutive data points reaching the 0.05 level of significance.

**Fig 3.**
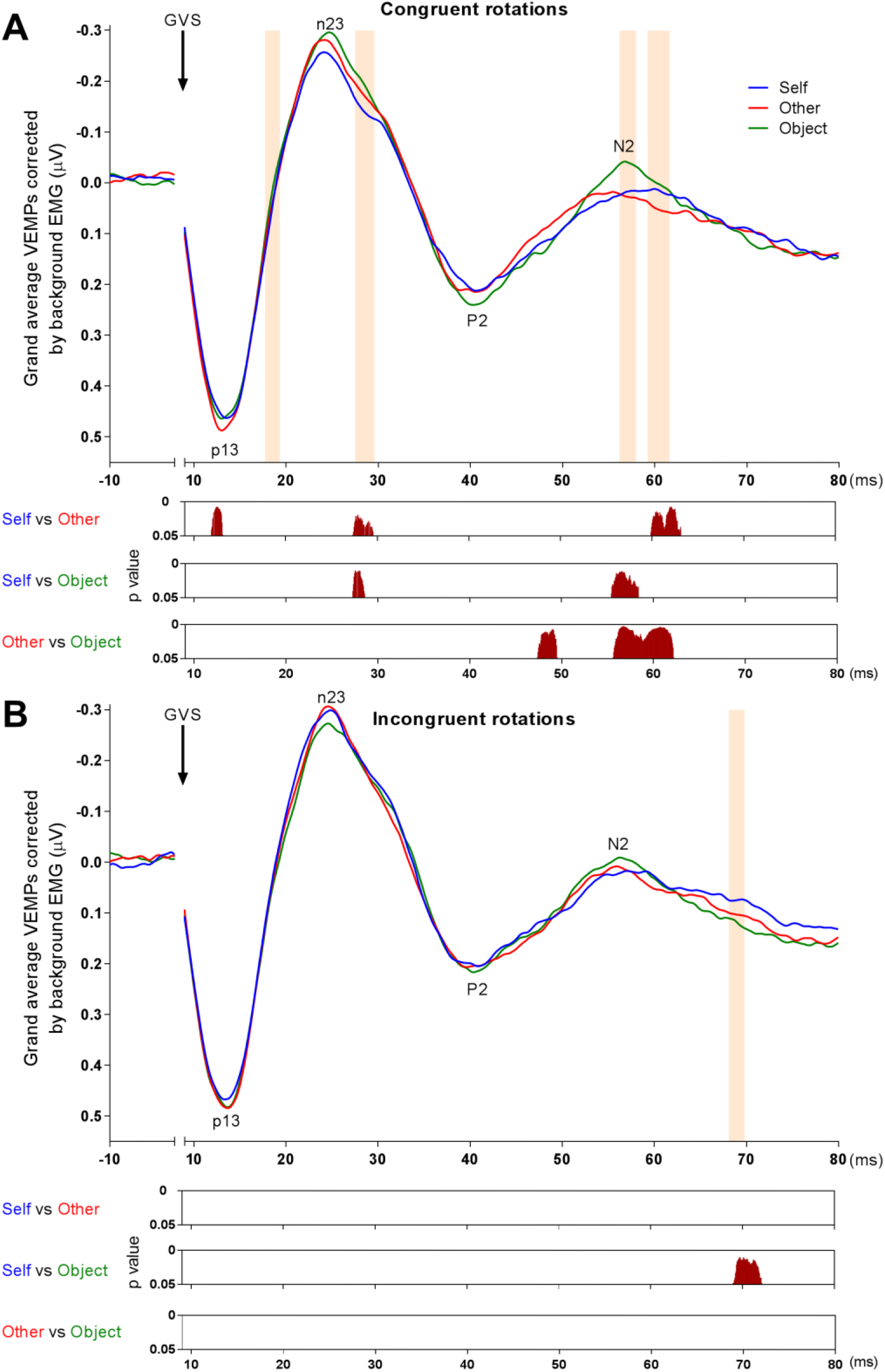
VEMPs recorded over the TRP muscle. Grand-average VEMPs (n = 20) are showed separately for Congruent rotations (A) and Incongruent rotations (B) for each category of Videos. Same conventions as for Figure 2.

We explored the main effect of Videos using Wilcoxon signed-rank tests (**Figures 2** and **3**). Overall, the analysis shows an attenuation of the VEMP during the observation of Self videos when compared to the observation of Other videos, revealing a modulation of the excitability of vestibulocollic reflexes by the person depicted in the video. This attenuation of the VEMPs was found for segments of the waveform spanning the p13-n23, as well as for the P2 and N2. When compared to Other videos, the amplitude of the peak of the p13 for Self videos was significantly reduced for both the SCM muscle (57 csdp., 11.5–15.0 ms) and TRP muscle (21 csdp., 11.9–13.1 ms). Similarly, the amplitude of the portion after the peak of the n23 was significantly reduced for Self videos when compared to Other videos for both SCM muscle (66 csdp., 22.4–26.4 ms, and 44 csdp., 27.2–29.8 ms) and TRP muscle (37 csdp., 27.3–29.5 ms). The amplitude of the N2 on the TRP muscle was also reduced during Self videos when compared to Other videos (54 csdp., 59.9–63.1 ms).

When compared to Object videos, observation of Self videos showed a significantly reduced p13 amplitude for the SCM muscle (49 csdp., 13.5–16.4 ms), as well as significantly reduced portion after the peak of the n23 for both muscles (SCM: 68 csdp., 23.2–27.3 ms; TRP: 22 csdp., 27.3–28.6 ms). The P2 amplitude was also significantly reduced for Self videos when compared to Object videos (SCM: 138 csdp., 40.0–48.3 ms) and this was also the case of the N2 amplitude (SCM: 46 csdp., 56.0–58.7 ms; TRP: 49 csdp., 55.4–58.3 ms).

The amplitude of the p13-n23 biphasic wave did not differ between the Other videos and the Object videos, suggesting that the attenuation of the early component reported above involves specifically the observation of self-motion. We note that the amplitude of the later components P2 and N2 recorded over both muscles was reduced for the Other videos when compared to the Object videos (P2 component: SCM, 93 csdp., 43.4–49.1 ms; TRP, 36 csdp., 47.4–49.5 ms and N2 component: SCM, 22 csdp., 55.4–56.7 ms; TRP, 109 csdp., 55.6–62.2 ms).

### Control for background EMG activity

There was no main effect of the Videos on the background EMG (**Figure 4**) on the SCM muscle (Friedman’s ANOVA, Congruent rotations: χ^2^(2) = 5.58, p = 0.61; Incongruent rotations: χ^2^(2) = 1.0, p = 0.61) nor on the TRP muscle (Congruent rotations: χ^2^(2) = 3.44, p = 0.18; Incongruent rotations: χ^2^(2) = 2.24, p = 0.33).

**Fig 4.**
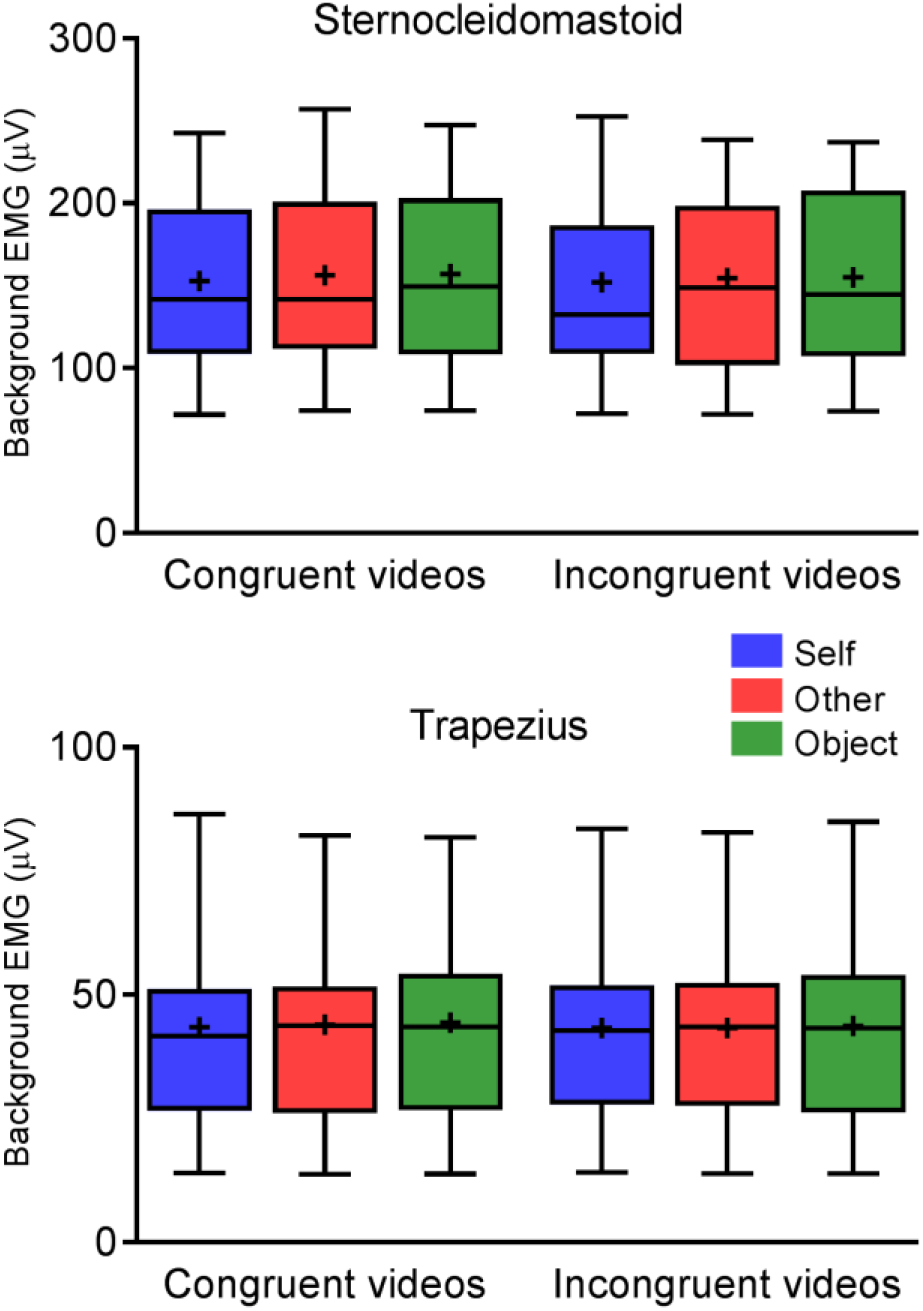
Background EMG activity. Box-and-Whisker plots illustrate the background EMG activity calculated over 6-s periods of video observation, for both SCM (upper panel) and TRP (bottom panel) muscles. The top and bottom ends of the whisker represent the 90^th^ and 10^th^ percentiles of the distribution, the bold horizontal line inside the box represents the median, the black cross represents the mean.

### Correlations between VEMP amplitude and empathy scores

The individual peak-to-peak p13-n23 amplitude on the SCM muscle correlated positively with the *empathic concern* scale for all Videos and both Directions of rotation (Kendall’s tau test, τ ≥ 0.40, all p ≤ 0.006) (**Table 1**). Thus, participants with higher empathic concern were those who tend to have larger vestibulocollic responses, irrespective of the video presented.

**Table 1.**
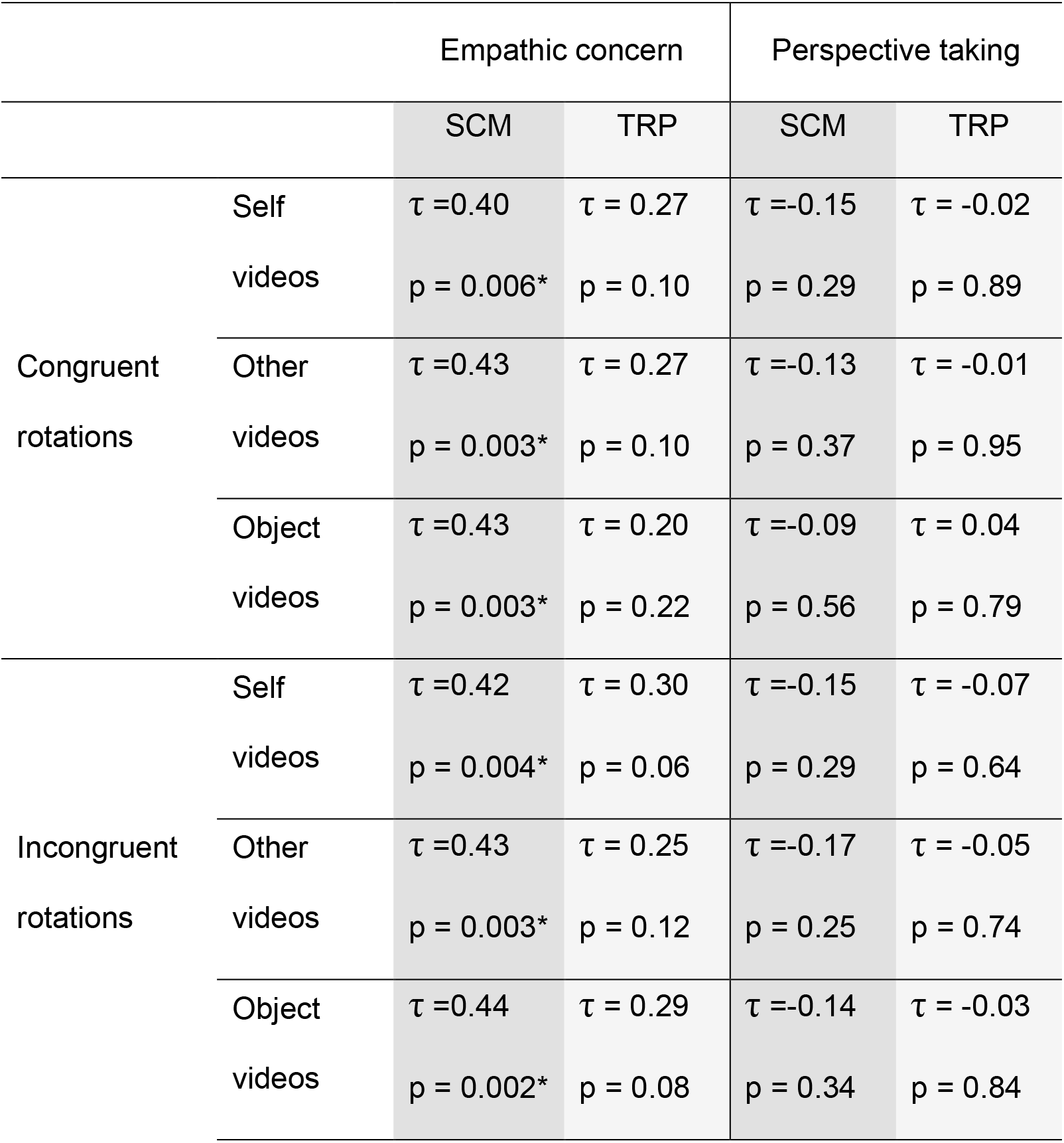
Correlation between individual peak-to-peak p13-n23 amplitude and scores to the *empathic concern* and *perspective taking* subscales of the IRI (Kendall’s tau test). Significant results at p < 0.05 are highlighted with an asterisk.

## Discussion

Our results provide original neurophysiological evidence that the excitability of vestibulocollic reflexes is sensitive to observing human bodies in motion, establishing connections between the so far distinct fields of social neuroscience and vestibular neurophysiology^22^. Although previous studies reported that attention^23^, motor imagery^24^ and emotions^12,25,26^ can influence the amplitude of vestibulo-spinal, vestibulocollic and oculomotor reflexes, the tasks and stimuli used in previous studies were devoid of self- and other-related visual information.

Observing the passive rotation of the self reduced the p13-n23, compared to observing the same rotation of another body or an object. However, observing the rotation of another body did not modulate the p13-n23 significantly, compared to an object. As for the somatosensory system^27,28^, it seems that there is something special about viewing one’s own body that modulates vestibular information processing.

Self information processing and self awareness has been associated with the insula^29,30^, with the precuneus, posterior cingulate cortex, temporoparietal junction, and the medial and anterior prefrontal cortex^31^. These areas overlap with several core regions of the vestibular cortex, such as the operculo-insular complex, temporoparietal junction and cingulate cortex^32,33^. Observing videos of the self may activate areas involved in both self-representations and vestibular information processing. We note that participants may also have recalled previous experience of vestibular sensations during the rotation of their own body. Such recall of vestibular sensations has been associated with bilateral activations of the inferior frontal gyri, anterior operculum, middle cingulate cortex, premotor cortex and anterior insula^34^. By activating cortical areas involved in self-representations, vestibular information processing and recall of self-motion, observing Self videos may have modulated VEMPs waveform through direct or indirect projections from the cortex to the vestibular nuclei^35,36^. Indeed, electrical stimulation of the multisensory vestibular cortex have been shown to activate or inhibit responses in the cat vestibular nuclei and these stimulations influenced vestibulo-spinal, vestibulocollic and oculomotor reflexes^37,38^.

Furthermore, electrophysiological recordings in monkeys showed that active head rotation strongly decreases the firing rate of vestibular nuclei neurons, compared to passive self-motion^39,40^. We propose that observing Self videos (while maintaining the head rotated towards the anode) triggers partly similar neurophysiological mechanisms to active self-motion and decreases the excitability of vestibulocollic reflexes. Indeed, we found a trend for stronger illusory self-motion for the observation of Self videos, which could be misinterpreted as active self-rotation.

Finally, we found that observing the passive rotation of the self also reduced the P2 and N2 responses, compared to observing another body (SCM) and an object (SCM and TRP). There is a controversy as to whether components following the n23 are of vestibular origin. Most authors proposed that later components are not of vestibular origin, as they survive after vestibular neurectomy^41^, whereas others reported opposite findings for the TRP muscles^42^. As most clinical and theoretical studies have disregarded those later components, it is unknown how P2 and N2 can be modulated by cognitive and emotional factors. The P2 and N2 are likely of multisensory origin, including vestibular, cochlear and somatosensory origins^42^. The modulation of the P2 and N2 components by low level information about self and other bodies is a new finding. It may reflect more complex multisensory mechanisms related to self-other resonances, i.e. the modulation of sensorimotor processing when observing bodies.

Just as observing other bodies in motion can have a detrimental effect on the observer’s balance^16,43^ and self-motion perception^17^, we found that observing another body undergoing passive motion reduced VEMPs waveform when compared to observing an object in motion. Surprisingly, the VEMP attenuation was found for components P2 and N2, but not for the early p13-n23 component.

The fact that observing both Self and Other videos decreased the P2-N2 amplitude – but to a lower extent for the conspecifics – indicates a modulation of the late VEMP components by self-other representations. This suggests that sensorimotor resonance between self and others also applies to the vestibular system. Sensorimotor resonance refers to the fact that observing another person receiving a sensory stimulation changes our processing of the same stimulation. It has been described extensively for other senses than the vestibular system and relies on common structures processing self and other sensory information^44–47^. Observing bodies undergoing passive whole-body motion may activate self and other representations and self-other resonance, modulating activity in the vestibular nuclei and decreasing the excitability of vestibulocollic reflexes through corticofugal projections.

Interestingly, the VEMPs waveform was only modulated by videos presenting Congruent rotations. First, this direction-specific effect rules out non-specific emotional and attentional modulation of the excitability of vestibulocollic reflexes. Second, it suggests that a specular congruency between the actual head position on the trunk and the direction of the observed rotation is more likely to influence the excitability of vestibulocollic reflexes. This is consistent with behavioral data suggesting that sensorimotor resonance becomes more important when the observed body posture or movement is compatible with the observer’s motor stabilization strategy^48^ and that third-person perspective taking is facilitated when the observer and the seen conspecific share a common body posture^49^. As vestibular nuclei neurons are sensitive to the position of the head on the trunk^50^, our data suggest that the specular congruency between visual and neck somatosensory signals facilitates the inhibition of vestibular nuclei neurons projecting to the spinal cord.

We found that the p13-n23 amplitude on the SCM, the muscles most strongly involved in the head rotation towards the cathode, was positively correlated with *empathic concern,* for all categories of videos. This suggests a general effect of empathy on the excitability of vestibulocollic reflexes, irrespective of the person observed. A recent functional magnetic resonance imaging study has linked the strength of visuo-vestibular responses to personality traits, with stronger responses in the vestibular nuclei and parieto-insular cortex of participants with higher neuroticism (i.e. more nervous participants) and larger responses in the amygdala of more introverted individuals^21,51^. Our data are in line with such a general effect of personality traits on vestibular information processing, extending previously defined interplay between emotional and social neural networks with the vestibular neural network^52–54^. A study^17^ revealed that empathy scores positively correlated with the congruent vs. incongruent latency difference to detect passive self-motion when simultaneously observing Others or Objects videos, but not Self videos. This suggests that personality traits impact multisensory self-motion perception (based on visual and vestibular signals) and vestibulocollic reflexes with different effects depending on the person depicted in the video. Thus, self-motion perception may involve more fine-grained multisensory – and social – regulation mechanisms^39^ than the excitability of vestibulocollic reflexes.

Our results show that vestibular information processing is sensitive to observing human bodies in motion, establishing new connections between research in social and vestibular neuroscience. From an evolutionary perspective, the present findings suggest that the human balance system evolved to react differently to moving objects and conspecifics. Similar studies in non-human primates could put our results in a comparative perspective and provide detailed information about the neurophysiological mechanisms involved. We note that previous studies of sensory processing in a social context have manipulated higher-level features of self-other resonance, such as political ideas, ethnicity, or pleasantness of the others^55,56^. While this was beyond the scope of the present investigation, our results may lead the way to the study of self-motion perception and vestibular information processing as a function of such social features. Finally, studies of the interplay between self-other representations, higher-level social features and vestibular information processing may have important applications for understanding balance disorders and improving their rehabilitation.

## Methods

### Participants

Twenty-five healthy volunteers participated (13 females; mean age ± SD: 23 ± 3 years), of whom 24 participants were right-handed (mean laterality quotient ± SD: 83 ± 16 %; Edinburgh Handedness inventory^57^) and one was left-handed (−40 %). They had normal or corrected-to-normal vision and declared no history of vestibular, neurological, or psychiatric disease. All participants provided written informed consent prior to participation. Experimental procedures were approved by the local Ethics Committee (Comité de Protection des Personnes Sud-Méditerranée II, 2011-A01221-40) and followed the ethical recommendations laid down in the Declaration of Helsinki.

### Visual stimuli

Visual stimuli consisted of videos showing the passive rotation of the participant (“Self videos”), of an unknown, age- and gender-matched person (“Other videos”), or of a white rectangular cuboid (“Object videos”), installed on the same rotating chair (Robulab 80, Robosoft SA, Bidart, France) (**Figure 1A**). Self-videos were recorded before electrophysiology recordings. Participants were seated on a rotating chair with their hands on their laps, eyes closed and a neutral face. A video camera (Sony HDR-XR160, Sony, Surrey, United Kingdom) placed 1.65 m in front of the participants recorded the rotation of their entire body on the chair. A black fabric behind the chair excluded all visual references from the background. Participants were rotated passively in the clockwise and counterclockwise direction around their longitudinal axis during 6 s with a sinusoidal velocity profile and a peak velocity of 18 °/s. The Other videos were recorded using the same procedures and with the same motion profile. An actor and an actress, who did not take part in the electrophysiology experiment, helped to create the videos depicting an unknown male and female body. The unknown body shown in the video was age-matched because our participants were all within the same age range. Object-videos were created following the same procedures, i.e. by rotating the white rectangular cuboid placed on the rotating chair. The rectangular cuboid was made of cardboard and had about the same height (84×31×31 cm) as the participants when seated on the chair. All videos were edited to last 6 s and were cropped to display the participant/actor from their head to their lower legs (when facing the camera) using Adobe Premiere Pro CC 2015. Participants were first seen from the side, and at the middle of the rotation (3 s), they were facing the camera (**Figure 1A**). During the experiment, videos were showed in a head-mounted display with a 30° horizontal field-of-view (LDI-100BE, Sony). This allowed us to maintain visual stimuli fixed in a head-centered coordinate system.

### Galvanic vestibular stimulation

Transmastoid galvanic vestibular stimulation (GVS) was used to evoke VEMPs recorded over two neck muscles^8,58^. A pair of carbon and rubber electrodes (4 × 2.5 cm, Plate electrode EF 10, Physiomed Electromedizin AG, Schnaittach, Germany) inserted in wet sponges was maintained on the skin covering the opposite mastoid processes using a cohesive contention strip around the head. Thirteen participants (7 females) had left cathodal/right anodal GVS configuration, whereas 12 participants (6 females) had right cathodal/left anodal GVS configuration. GVS consisted in series of square-wave pulses of 2 ms delivered at 8 Hz with an intensity of 3.2 to 5.0 mA (depending of the participant’s skin sensitivity; mean intensity ± SD:4.7 ± 0.7 mA) by a Grass S88 stimulator (Grass Instrument Co, Quincy, Massachusetts, USA) coupled to a constant current and isolating unit (Grass PSIU-6B). Short-duration GVS does not evoke self-motion perception. GVS was preferred over acoustic air-conducted stimulation of otolithic receptors as it provides more physiologically valid inputs related to bilateral stimulation of the vestibular receptors. GVS increases the firing rate in the vestibular afferents under the cathode, while decreasing the firing rate in the afferents under the anode^59^.

### Electromyography recordings

VEMPs were recorded over the sternocleidomastoid (SCM) and trapezius (TRP) muscles following previously described procedures^8,12^. Active electrodes (FLAT Active electrode, Biosemi Inc., Amsterdam, Netherlands) were placed at the junction of the upper and middle thirds of the SCM ipsilateral to the cathode and at the intersection between the upper and middle muscle fibers on the TRP ipsilateral to the cathode. In our system, the Common Mode Sense (CMS) and Driven Right Leg (DRL) electrodes replace the single standard ground electrode and form a feedback loop to increase the signal-to-noise ratio. CMS-DRL electrodes were placed 2 cm apart over the C7 vertebra. A reference electrode was placed on the sternum. Preamplified electromyographic signals (EMG) were sampled at 16 kHz with a bandwidth of 0.16−3200 Hz and analyzed offline using custom-made scripts in Matlab R2015b (The MathWorks Inc., Natick, USA).

As cervical VEMPs are inhibitory responses of the ipsilateral neck muscles, participants were required to maintain tonic activation of the SCM and TRP ipsilateral to the cathode. They sat on a chair whose backrest was tilted ~45° backward. Participants actively maintained their head flexed towards the torso and rotated it ~80° towards the anode (the amplitude of head rotation was adapted to each participant to be as comfortable as possible). This contracted the SCM, and to a lower extent the TRP, under the cathode (**Figure 1A**). Participants were trained to maintain a stable muscular contraction before the experiment. In addition, the experimenter controlled the participant’s head position and level of muscle contraction on the visual display of the recording software (Actiview 7.03, Biosemi Inc., Amsterdam, Holland) throughout the experiment.

### Convention for the direction of rotation of the body/object in the videos

The direction of rotation of the body/object in the video was not referred to as clockwise and counterclockwise rotation, as there was no specific hypothesis about differences between those directions. By contrast, vestibular perception depends on the congruency between the direction of rotation of the body/object in the video and the actual direction of rotation of the observer’s body^17^. As VEMPs were recorded with the participant’s head maintained rotated towards the anode, we defined the direction of rotation of the body/object in the video according to the congruency of the observed rotation with the initial rotation of the participants’ head (**Figure 1B**). According to our convention, videos with “Congruent rotations” showed rotations in the direction matching what participants would have seen of their initial head rotation in a mirror, whereas videos with “Incongruent rotations” showed rotations in the opposite direction. Thus, Congruent rotations were rotations towards the anode, whereas Incongruent rotations were rotations towards the cathode.

### Experimental procedures

Each category of video (Self, Other, and Object) was presented 24 times for Congruent rotations, and 24 times for Incongruent rotations, resulting in a total of 144 visual stimuli per participant. Visual stimuli were presented in a randomized order in 18 blocks of 8 videos. Each block of visual stimuli started with the presentation of a white fixation cross on a black background for 500 ms, followed by a video. After each video a fixation cross was presented for 500 ms plus the response time to the detection task described below (with a maximum of 1 s). This resulted in a maximal duration of 56.5 s per block. Participants were asked to fixate on the cross presented between videos and to fixate on the body/object at the center of the screen during the videos. GVS pulses began with the video onset and were applied at 8 Hz during 6 s. Thus, 48 GVS pulses were presented during a video, resulting in a total of 1152 vestibular stimulation per category of video and direction of rotation. This high number of stimulation, compared to previous electrophysiological studies, ensured a good signal-to-noise ratio. Video presentation and GVS application were controlled by Superlab 4.5 (Cedrus Corporation, San Pedro, USA). Participants maintained a stable muscle contraction during each block of visual stimuli and rested as long as they wanted to relax their neck between consecutive blocks of visual stimuli.

To maintain their alertness during the recordings, participants were involved in a two-alternative forced choice task. In 25% of the videos, images were inverted vertically for 100 ms, starting at 1.5 s, 3 s or 4.5 s after the video onset (**Figure 1C**). Participants were asked to observe the body/object being rotated and to indicate as quickly and accurately as possible whether the video was, or was not, temporarily presented upside-down. They were instructed to answer during the fixation cross following the video, and we confirmed that no answer was given during the videos. Participants responded on two buttons of a response pad (RB-830, Cedrus Corporation, San Pedro, USA) with their right middle and index fingers (13 subjects responded ‘yes’ with their middle finger, 12 responded ‘yes’ with their index finger). Before the recordings, participants trained to the task on 10 trials without GVS.

### Illusory self-motion questionnaire

At the end of the experiment, participants filled out a questionnaire about illusory self-motion. For both directions of rotation (Congruent, Incongruent), participants answered the question “Did you feel a sensation of motion of your own body when observing the videos of yourself/the other person/the object?” Answers were given on a 7-point Likert scale ranging from “not at all” to “absolutely felt something”. Participants could indicate whether the illusory motion was in the same direction as the observed motion and add comments. Participants answered this questionnaire once about their average experience of the whole electrophysiology experiment.

### Interpersonal reactivity index

Participants completed the Interpersonal Reactivity Index^20,60^. Our analyses focused on the relation between VEMP amplitude and two subscales of the IRI measuring self-reported empathic concern and perspective taking. Both scales have been showed to correlate with implicit perspective taking^61^. The *empathic concern scale* includes seven questions assessing “other-oriented feelings of sympathy and concern for unfortunate others”, while the *perspective taking scale* includes seven questions assessing “the tendency to spontaneously adopt the psychological point of view of others”^20^. Ratings were completed on a 5-point scale ranging from “describes me very well” to “does not represent me very well”.

### Data recording and analysis

EMG signals were referenced to the electrode placed on the sternum and band-pass filtered (0.1−1000 Hz). All GVS pulses applied during the same category of videos (Self, Other, Object) and with the same direction of rotation (Congruent, Incongruent) were pooled together to calculate an average VEMP for each participant. VEMPs were calculated on epochs starting 25 ms before GVS pulses until 100 ms post-stimulus and were baseline-corrected by the average unrectified EMG during the 25 ms pre-stimulus. Epochs whose baseline (unrectified EMG) exceeded the mean ± 3 SD of the baseline were excluded, and the same was done for the rectified signal in the 100 ms post-stimulus. After data pre-processing, VEMPs on the SCM muscles were calculated on (mean ± SD): 1047 ± 101 GVS pulses for Self videos, 1049 ± 92 pulses for Other videos, and 1052 ± 92 pulses for Object videos (no effect of the category of videos, Friedman’s ANOVA: χ^2^(2) = 0.67, p = 0.72). VEMPs on the TRP muscles were calculated on 868 ± 65 GVS pulses for Self videos, 865 ± 71 pulses for Other videos, and 866 ± 66 pulses for Object videos (no effect of the category of videos, χ^2^(2) = 0.67, p = 0.72).

It is known that the level of background muscle activation is linearly correlated with VEMP amplitude^8,62^. Thus, we normalized each epoch by the average rectified EMG during the 25 ms pre-stimulus to account for the level of background EMG^12,63,64^. We compared SCM and TRP contraction for each category of videos and both rotations by calculating the average background EMG over the 25 ms pre-stimulus.

GVS evokes VEMPs with shorter latency^8^ than acoustic stimulation of otolithic receptors, as GVS bypasses the mechano-electrical transduction. Yet, we named “p13-n23” the first biphasic response on the SCM and TRP muscles in accordance with responses to acoustic stimulation^8^. In our study, the p13-n23 component was identified as the first salient positive-negative peak complex within a time window of 8 to 25 ms after GVS onset^65^. In our sample of participants, VEMPs were detectable over the SCM of 24 participants and over the TRP of 20 participants. Only their data were considered for subsequent analyses.

For each muscle, mean responses from these participants were averaged to obtain grand-average VEMPs for each category of videos and each direction of rotation. As most of the dependent variables were not normally distributed, we used non-parametric Friedman’s ANOVA and Wilcoxon signed-rank tests to conduct waveform analyses of grand-average VEMPs, as done for event-related potentials in electroencephalographic investigations of sensory and cognitive processing^66–68^. This approach allows a point-by-point analysis of the exact time course of the vestibulospinal reflex, without *a priori* hypotheses about the timing of the differences (only two points, the peaks of the p13 and n23, are classically analyzed). We corrected for temporal autocorrelation by using the constraints of 20 consecutive data points reaching the 0.05 level of significance^66,69^. Waveform analyses were conducted within 9 to 80 ms after GVS onset to span on waves of interest, that is on the p13 and n23 components, as well as later components (P2 and N2).

Finally, to analyze relations between VEMPs and empathy, we measured the peak-to-peak amplitude of the individual p13-n23 response^8,12^ and calculated the correlation coefficient of this amplitude with empathic concern and perspective taking scores.

## Acknowledgements

The research leading to these results has received funding from the People Programme (Marie Curie Actions) of the European Union’s Seventh Framework Programme (FP7/2007–2013) under REA Grant agreement number 333607 (‘BODILYSELF, vestibular and multisensory investigations of bodily self-consciousness’). The authors thank Ali Gharbi and Guy Escoffier for technical assistance, and Dr. Manuel Vidal for his help with the rotating chair.

## Author contributions

All authors participated to the experimental design. E.N. and D.D. recorded the data, E.N. analyzed the data. E.N., D.D. and C.L. contributed to statistical analysis. E.N. and C.L. wrote the main manuscript text. E.N. prepared Figs. 1–4 and Table 1. All authors reviewed the manuscript. Funding was provided by C.L.

## Additional information

The authors declare no competing interests.

## Notes

### Competing Interest Statement

The authors have declared no competing interest.

